# Differential signaling by two *Plasmodium* sporozoite adhesins mediates transmission of malaria parasites

**DOI:** 10.64898/2026.05.22.727197

**Authors:** Monami Roy Chowdhury, Marzia Matejcek, Dennis Klug, Bastian Löhning, Bea Jagodic, Jessica Kehrer, Franziska Hentzschel, Friedrich Frischknecht

**Affiliations:** Parasitology, Center for Infectious Diseases, Medical Faculty, Heidelberg University, 69120 Heidelberg, Germany; Department of Molecular Cell Physiology, Institute of Physiology and Pathophysiology, Philipps-University Marburg, Marburg, Germany; Center for Infection Research, DZIF, partner site Heidelberg, Germany; SynthImmune Cluster of Excellence, Heidelberg University, 69120 Heidelberg, Germany

**Keywords:** gliding motility, invasion, Plasmodium, malaria, transmission, adhesins

## Abstract

Malaria parasites are transmitted in the form of *Plasmodium* sporozoites, highly motile cells that form in oocysts at the mosquito midgut and invade salivary glands. After transmission to the vertebrate host, sporozoites migrate in the skin and liver to enter and differentiate in hepatocytes. Sporozoite migration and invasion are mediated by surface adhesins which link the extracellular substrate to the actin-myosin motor, providing the force for gliding motility. Here we show that two adhesins, TRAP and TRP1, use different extracellular domains and distinct cytoplasmic signaling to mediate salivary gland invasion and gliding motility. The extracellular thrombospondin repeat (TSR) of TRP1 but not the TSR of TRAP is essential for gliding and transmission and the TSR of TRAP can only partially rescue TSR function in TRP1. Furthermore, the cytoplasmic domain of TRAP cannot replace the equivalent domain of TRP1, which is essential for gliding and salivary gland invasion. This suggests that sporozoites integrate signals through multiple essential outside-in signaling pathways to achieve migration and invasion.

## Introduction

Migration and invasion are tightly coupled processes in apicomplexan parasites and essential for the propagation of their life cycles (1). Both processes are mediated by receptor-ligand interactions that signal across the plasma membrane of the parasites and by an actin-myosin motor that propels the parasite forward. *Plasmodium* sporozoites are the forms of the malaria parasite transmitted from mosquitoes to humans or other animals (2). Sporozoites form in and egress from oocysts located at the midgut wall of the mosquito, penetrate salivary glands to accumulate in the salivary cavities and are deposited in the dermis of the host during the probing phase of a mosquito bite. In the dermis, sporozoites migrate at high speed to find and enter blood vessels. Sporozoites travel with the flow of the blood and extravasate in the liver to specifically invade hepatocytes and differentiate into red blood cell-invading merozoites (2, 3). Sporozoite migration is driven by an actin-myosin motor coupled to adhesins of the thrombospondin related anonymous protein (TRAP) family (3–5). Actin filaments are polymerized by formin 1 anchored to the apical tip of the parasite (6–8), and TRAP family adhesins are secreted to the surface at the front end of the parasite by micronemes, small vesicles (4). Upon secretion, TRAP family adhesins are transported rapidly to the proximal end of the sporozoite by the actin-myosin motor (4, 9). TRAP family adhesins are thought to engage with unknown ligands on the surface of cells or non-cellular substrates using their integrin-like I domain (10). Upon ligand engagement, a conformational change of the I-domain from the closed to the open form leads to outside-in signaling, possibly resulting in the engagement of the TRAP family adhesin to actin filaments (11, 12). TRAP is the most strongly expressed member of TRAP family adhesins in sporozoites (13). Next to the I-domain this protein also contains a thrombospondin repeat (TSR) and a cytoplasmic domain that features a penultimate tryptophan. Deletion of *trap* allows sporozoites to escape from the oocysts but these *trap*(*-*) sporozoites cannot enter the salivary gland (3). Other TRAP family proteins contribute to the efficiency of salivary gland invasion or migration in the skin (5). The thrombospondin related protein 1, TRP1, lacks an I domain and does not show a penultimate tryptophan. Deletion of *trp1* leads to sporozoites that are trapped inside oocysts (14). Hence, both TRAP and TRP1 are essential for transmission from mosquito to mammals (Figure 1A). Both adhesins are proteolytically processed, TRAP by a membrane-resident rhomboid protease (15) and TRP1 by an unknown protease (14). Notably, *trp1*(-) sporozoites are not motile in oocysts and fail to egress while *trap*(-) sporozoite can readily egress from oocysts, suggesting that *in vivo* TRP1 might mediate the activation of motility ‘upstream’ of TRAP. Curiously, when isolated from oocysts and placed in motility-activating medium *in vitro trp1*(-) sporozoites can move (14). We thus hypothesize that once sporozoites are outside oocysts TRAP alone is sufficient for conferring motility. This observation suggests the existence of two different signaling routes to activate migration *in vivo* through TRP1 and TRAP. Here we probe this hypothesis by generating and analyzing a number of transgenic parasite lines expressing TRP1 with targeted mutations and deletions in the TSR, deletion and swaps of the cytoplasmic domain as well as parasites expressing TRAP lacking the TSR. Our data shows that the TSR of TRAP is largely dispensable, while the TSR of TRP1 is essential for TRP1 function. Replacing the C-terminus of TRP1 with that of TRAP does not rescue TRP1 function suggesting a differential outside-in signaling for sporozoite migration and invasion through these two adhesins.

**Figure 1.**
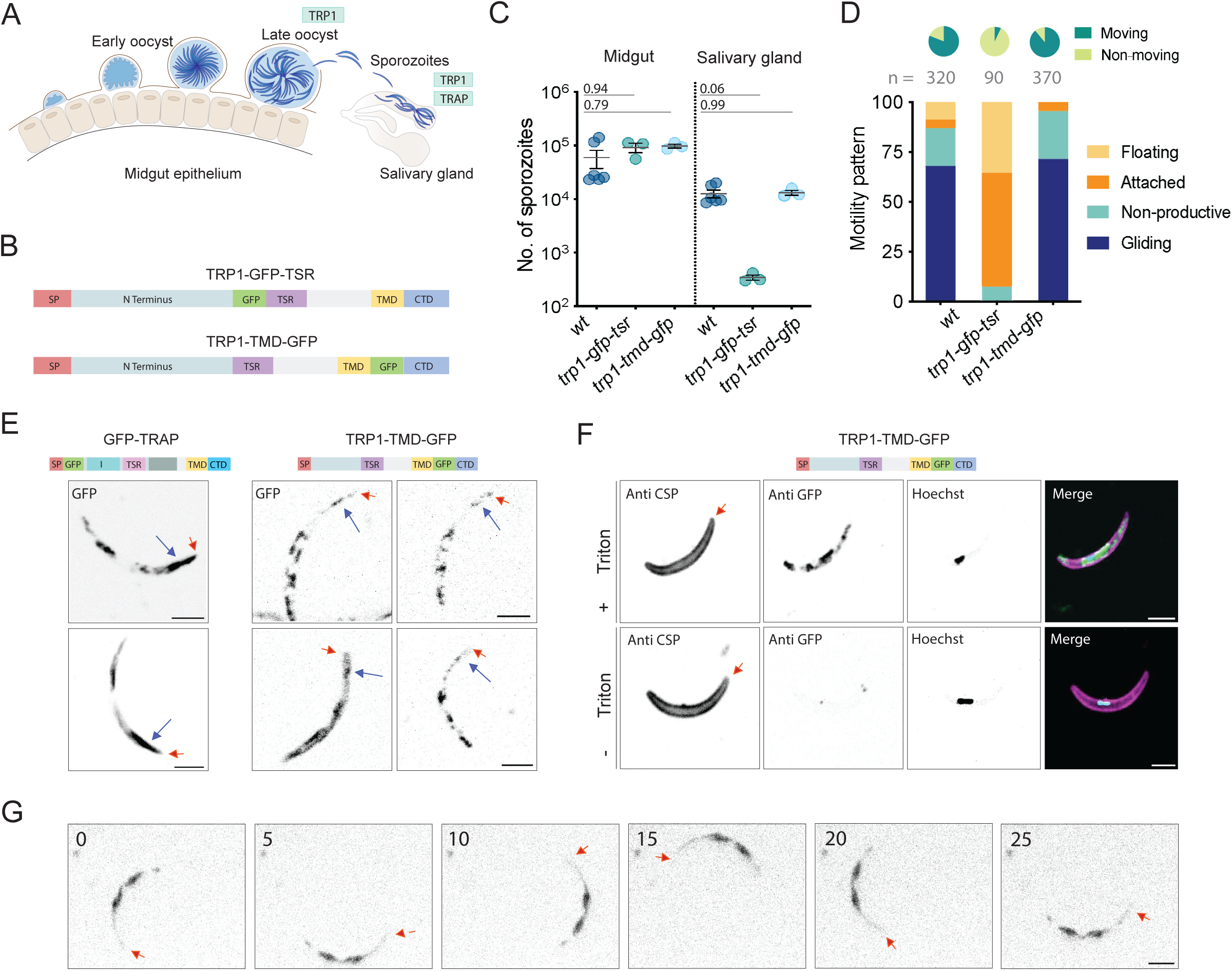
TRP1 localizes in stable patches on the sporozoite periphery. A. Sporozoite development in *Plasmodium* oocysts at the midgut wall and invasion of salivary glands. Note that TRP1 is important for egress from oocysts and invasion of salivary glands, while TRAP is important for invasion of salivary glands only. B. Schematic representation of TRP1-GFP-TSR and TRP1-TMD-GFP. C. Total numbers of sporozoites per midgut and salivary gland of wild type (wt) and TRP1-GFP-tagged lines; each dot represents an average number of sporozoites calculated from mosquitoes dissected from an independent cage. Black lines represent mean while the error bars represent SEM. D. Motility patterns of salivary gland sporozoites observed in wild type (wt) and both TRP1 GFP-tagged lines. ‘n’ denotes the total number of sporozoites quantified from each parasite line. The pie charts depict the proportion of moving and non-moving sporozoites overall in each parasite line, where both productive and unproductive motility accounts for the ‘moving’ sporozoites. Attached and floating sporozoites were clubbed together into the ‘non-moving’ counterpart. E. Localization of GFP in living *gfp-trap* and *trp1-tmd-gfp* salivary gland sporozoites, scale bar: 3 μm. Red arrows point to the front end of the parasite. Long blue arrows point to micronemal localization. F. Localization of GFP in *trp1-tmd-gfp* salivary gland sporozoites in the presence or absence of Triton X-100, indicating a patchy localization throughout the sporozoites (scale bar: 3 μm). Red arrows point to the front end of the parasite. G. Time lapse images of *trp1-tmd-gfp* salivary gland sporozoites indicating that the GFP signal is not dynamic in moving sporozoites. Red arrows point to the front end of the parasite. Statistical significance calculated by Kruskal-Wallis with Dunn’s multiple comparisons test (Data collected from at least three independent cage feed experiments).

## Results

### TRP1 localizes in stable patches on the sporozoite periphery

TRAP resides in micronemes from where it is secreted to the surface (16, 17). To investigate TRP1 localization we previously attempted endogenous functional tagging at the N-terminus and C-terminus, which however failed to yield fully infectious sporozoites (14). Therefore, we here used internal GFP fusions, in analogy to our recent work with circumsporozoite protein (CSP) (18, 19). We generated two *P. berghei* strain ANKA parasite lines (Supplementary Fig. 1) with GFP inserted either N-terminally of the TSR (*trp1-gfp-tsr*) or C-terminally to the transmembrane domain (*trp1-tmd-gfp*) of the endogenous protein with a small linker (Figure 1B). While sporozoites of both lines were formed in normal numbers, only those with the GFP fused C-terminally to the transmembrane domain would efficiently enter salivary glands (Figure 1C). Movement analysis revealed that the TRP1-GFP-TSR salivary gland sporozoites did not migrate, while the *trp1-tmd-gfp* salivary gland sporozoites moved as well as wild type (Figure 1D). Salivary gland-derived *trp1-tmd-gfp* sporozoites showed a striking pattern of fluorescence. Unlike *gfp-trap* (20), the *trp1-tmd-gfp* signal was only weakly associated with micronemes and showed a strong patchy pattern near the surface, which appeared in a unique distribution for each sporozoite (Figure 1E). Treatment of *trp1-tmd-gfp* sporozoites with and without triton to permeabilize the plasma membrane followed by staining with an anti-GFP antibody revealed a signal only for triton treated parasites, suggesting that the GFP is, as predicted, localized inside of the parasite (Figure 1F). Imaging motile *trp1-tmd-gfp* sporozoites further showed that the fluorescent signals stayed in the same place with respect to the sporozoites (Figure 1F). These data show that TRP1 is strongly expressed in salivary gland sporozoites and localizes differently than TRAP.

### The TSR of TRP1 is essential for transmission

The observation that the insertion of a GFP near the TSR of TRP1 abrogated sporozoite motility (Figure 1C, D) gave rise to the hypothesis that the TSR is essential for migration. Yet, in TRAP, previous work using point mutations suggested only a minor role of the TSR for transmission of sporozoites (21). These studies focused either on introducing individual point mutations in the conserved tryptophan residues in TSR (21) or on removing only the TSR core segment (15 amino acids: PfW250–PfR264; PbW244–PbR258) in a chimeric *P. berghei* line engineered to express the TRAP of the human-infecting parasite *Plasmodium falciparum*, i.e. PfTRAP instead of PbTRAP (22). In contrast to the TSR domain, while point mutations in the I-domain allowed for some invasion of salivary glands, deletion of the entire domain nearly completely abrogated salivary gland invasion (10, 21). We thus tested if deletion of the TSR in TRP1 or TRAP would reveal a stronger phenotype as the one reported by targeted mutations.

To investigate a potential role of the TSR in TRP1, we generated a parasite line lacking this domain (*trp1Δtsr*) as well as two further parasite lines, one with point mutations targeting the conserved tryptophans (*trp1-tsr-AA*) and a second one featuring a complete domain swap with the TSR of TRAP (*trp1-tsr^TRAP^*) (Figure 2 A,B, Supplementary figure 2). Analysis of the lines revealed that, as expected, all lines formed oocysts similar to wild type (Table 1, Supplementary Figure 2,3). Interestingly, while all sporozoites egressed from oocysts normally, as indicated by comparable numbers of sporozoites in the mosquito hemolymph, the sporozoites of the *trp1Δtsr* line largely failed to enter salivary glands (Figure 2C). Notably, both the *trp1-tsr-AA* and the *trp1-tsr^TRAP^* mutants also showed a notable, albeit statistically non-significant reduction in salivary gland invasion. Analysis of the movement of sporozoites placed on glass showed that there was a strong reduction of productive motility in the TRP1-TSR domain mutants (Figure 2D). A reduction in salivary gland sporozoite numbers and motility usually goes along with a decrease in transmission of the parasites. To investigate this, we let ten infected mosquitoes bite a naïve mouse and monitored the emergence of blood stage parasites and the development of blood stage infection (parasitemia). This resulted in no transmission to host for all TRP1-TSR domain mutants. Intravenous (i.v.) injection of 1.000 salivary gland derived sporozoites resulted in a strong decrease in infectivity in TRP1-TSR domain mutants with at least two days delay in prepatency and some animals not becoming infected (Figure 2E).

**Figure 2.**
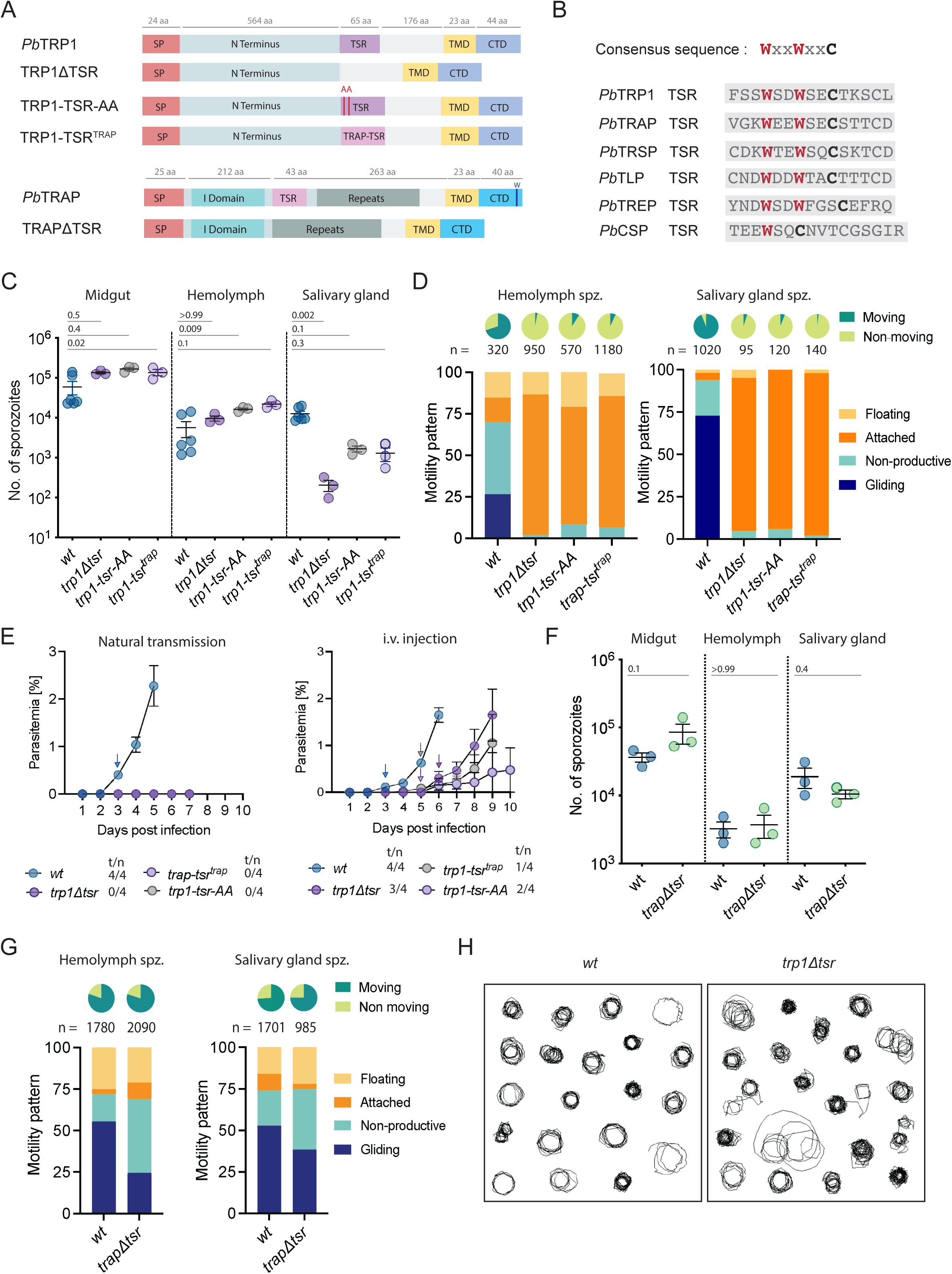
The TSR of TRP1 but not TRAP is essential for transmission. A. Schematic representation of *Pb*TRP1 and *Pb*TRAP along with the indicated domain mutants. B. Amino acid sequence alignment of TSR domain core across various proteins expressed in sporozoites, highlighting the conserved tryptophan residues. C. Total numbers of sporozoites per midgut, hemolymph and salivary gland of various TRP1 TSR domain mutants; each dot represents the average number of sporozoites calculated from mosquitoes dissected from one independent cage. Black lines represent mean while the error bars represent SEM. D. Motility patterns of hemolymph and salivary gland derived sporozoites from various TSR domain mutants and wild type. E. Natural transmission and intravenous injection (1000 salivary gland sporozoites per mouse) assay with the indicated parasite lines. t/n: number of mice showing blood stage infection / number of challenged mice. Black lines represent mean while the error bars represent SEM. F. Total numbers of sporozoites in midgut, hemolymph and salivary gland of the TRAP TSR domain deletion and wild type control; each dot represents the average number of sporozoites calculated from mosquitoes dissected from an independent cage. Black lines represent mean while the error bars represent SEM. G. Motility patterns of hemolymph and salivary gland derived sporozoites from the TRAP TSR domain deletion line and wild type controls. H. Tracks of 20 randomly selected sporozoites from the TRAP TSR domain deletion line and wild type controls. Statistical significance calculated by Kruskal-Wallis with Dunn’s multiple comparisons test (Data from three independent cage feeds).

**Table 1:**
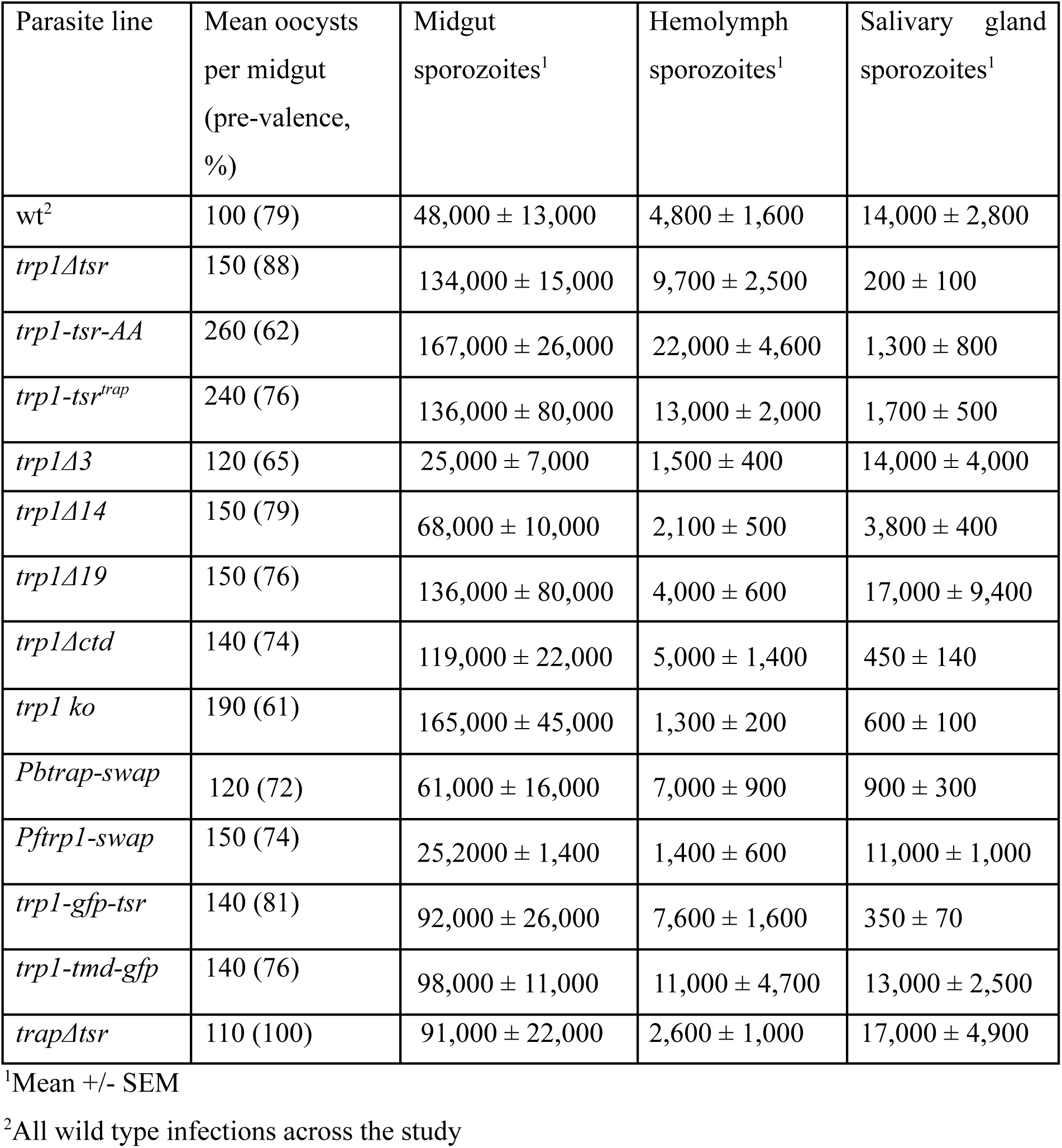
Average numbers of parasites in different tissues of all reported lines.

To explore the importance of the role of the TSR domain in TRAP, *P. berghei* parasites were generated that expressed a version of TRAP lacking the TSR (Figure 2A, Supplementary Figure 2). The *trapΔtsr* parasite line formed oocysts similar to wild type (Table 1, Supplementary Figure 2,3) and also entered salivary glands at levels comparable to wild type (Figure 2F). Analysis of the movement of sporozoites placed on glass showed that there was a slight decrease in gliding motility of the *trapΔtsr* line (Figure 2G). This was reflected by a slightly decreased sporozoite speed in persistently gliding sporozoites and a slightly less persistent circular motility (Figure 2H). Natural transmission assays and i.v. injection of 1,000 or 10,000 salivary gland-derived *trapΔtsr* sporozoites revealed that there was no delay in infectivity to the host in *trapΔtsr* mutants in natural infections and when 10,000 sporozoites are injected but a one-day delay in one experiment injecting 1,000 sporozoites (Table 2). These data suggest an essential role for the TSR domain in TRP1 in transmission of malaria parasites from mosquitoes to the mammalian host but a much weaker role for the TSR of TRAP.

**Table 2:**
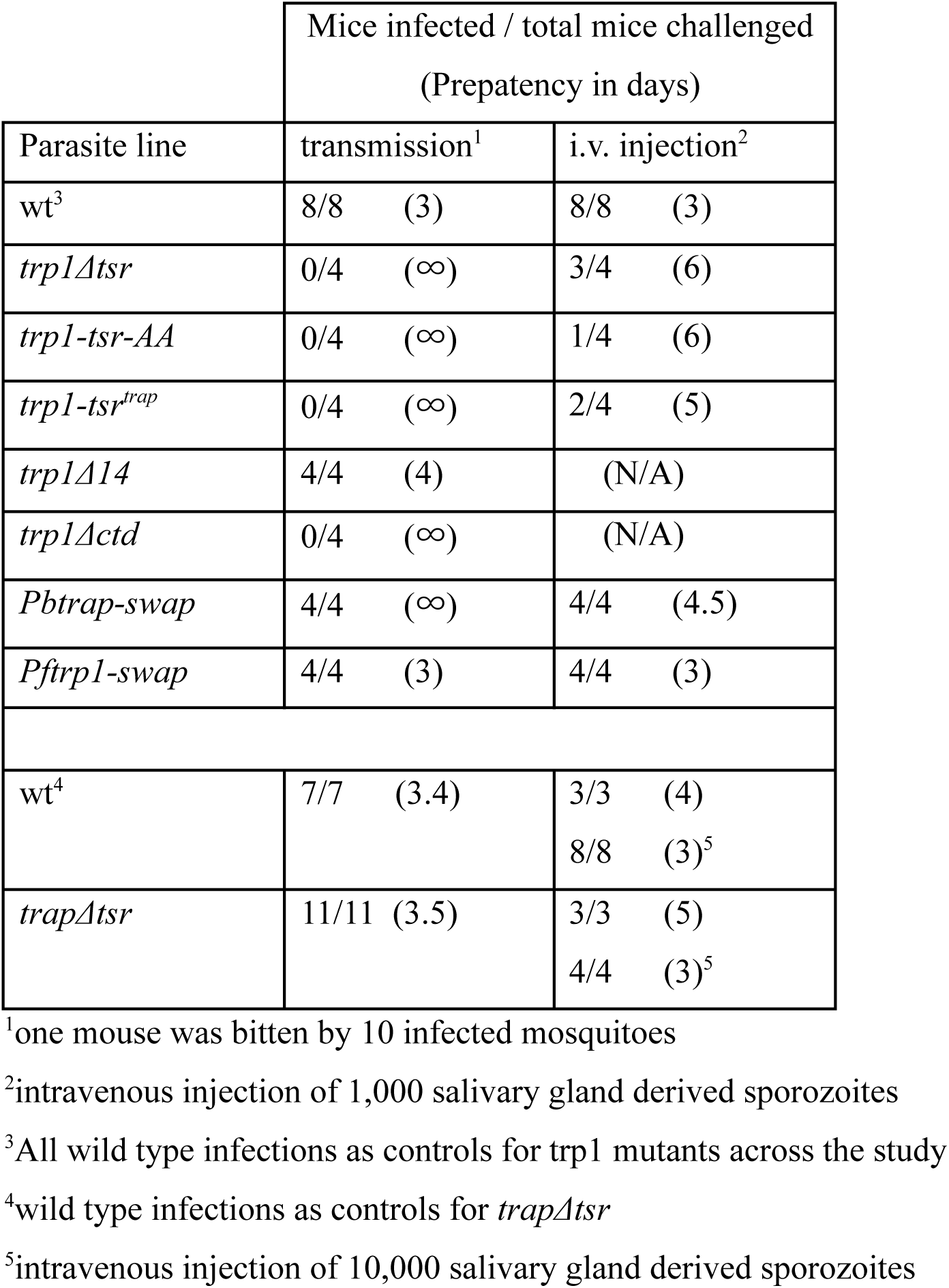
Transmission capacity of the reported lines.

### The C-terminus of TRP1 is essential for transmission

TRAP family adhesins interact via their C-terminus with proteins linking the adhesins to actin filaments (7). Deletion of the C-terminus of TRAP led to parasites that could not move productively and failed to enter salivary glands (23). Previous work based on the deletion of the entire C-terminus suggested that the C-terminus of TRP1 is also essential for invasion of salivary glands (14). To understand which part of the C-terminus is essential for the function of the protein, we generated three different parasite lines lacking the last 3, 14 or 19 amino acids (Figure 3A, Supplementary Figure 4). As expected, these mutations did not affect progression of the parasite into the mosquito (Supplementary Figure 4). Yet surprisingly, we found a decreased number of parasites in the salivary glands for the parasite lacking 14 amino acids, but not for those lacking 3 or 19 amino acids (Table 1, Figure 3B). Imaging of salivary gland-derived sporozoite revealed a striking expansion of a rare, and previously overlooked movement pattern for sporozoites (Figure 3C). Wild type parasites undergo mostly progressive circular gliding, while some also show a waving motion, where the parasite is attached at one end and actively moves the body within the medium (24). The parasite line lacking 3 amino acids showed similar movement patterns. In contrast, around half of the mutant sporozoites population lacking the last 14 C-terminal amino acids of TRP1 and some lacking 19 amino acids moved in an unusually erratic way (Figure 3C,D). Short circular movements were interspersed with back-and-forth movements and waving patterns (Figure 3C). We termed this new motility type “wave-flipping”. This different gliding pattern and the decreased number of sporozoites in the salivary glands suggest an important function of the C-terminus in salivary gland entry. To test for the impact on transmission of the parasite lacking 14 amino acids, we let infected mosquitoes feed on naïve mice and followed the onset and course of blood stage infection. This showed that all mice could be infected by the mutants, albeit at a delayed blood stage onset of 1 day, reflecting a decrease of 90% in infectivity (25) (Figure 3E). Together these data suggest that the C-terminus of TRP1 is important for salivary gland invasion and might be folded in a complex manner to interact with its partners.

**Figure 3.**
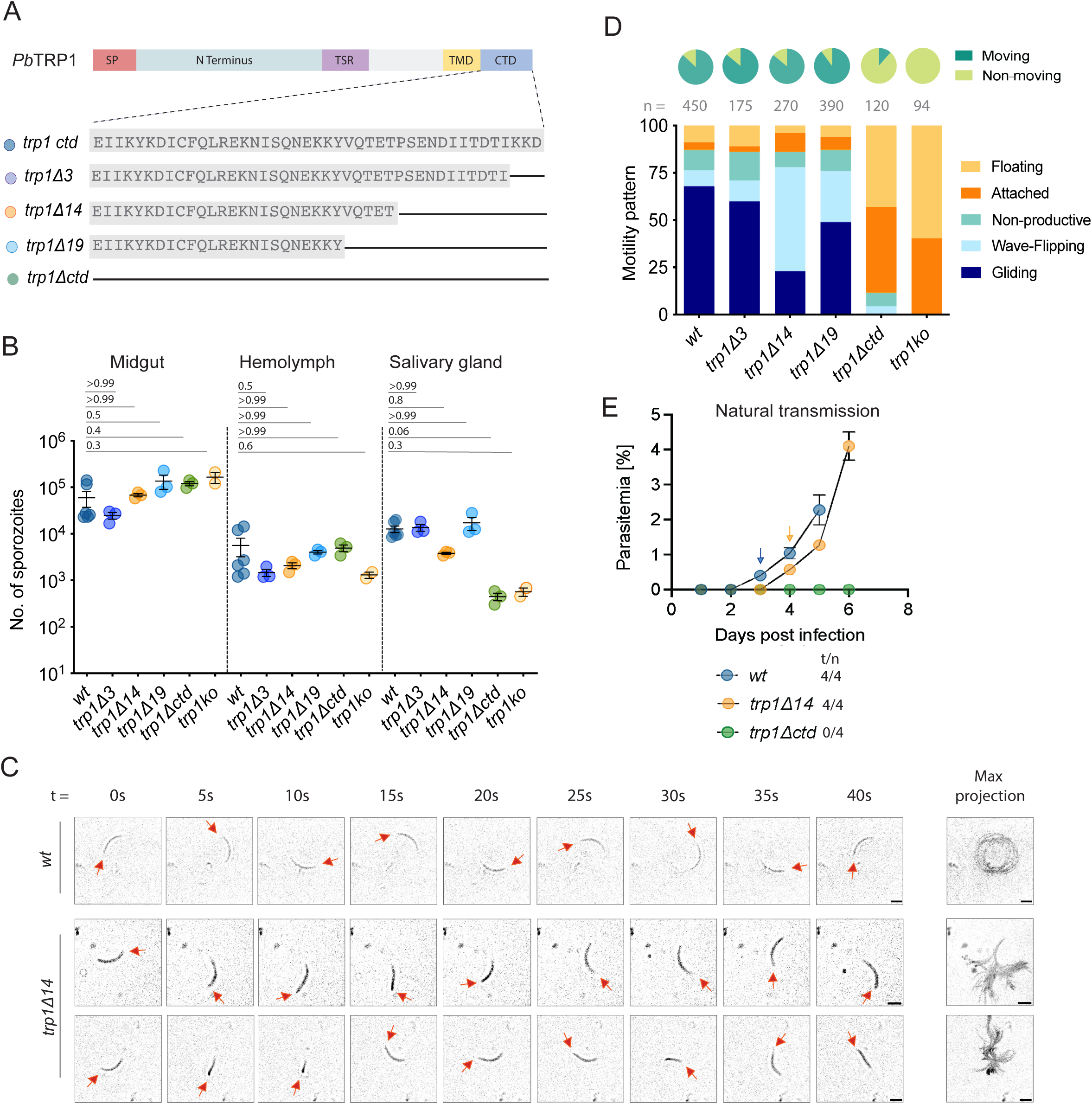
The C-terminus of TRP1 is essential for transmission. A. Schematic representation and amino acid sequences of TRP1 C-terminus domain deletion mutants. B. Total numbers of sporozoites per midgut, hemolymph and salivary gland of various C-terminal deletion mutants; each dot represents the average number of sporozoites calculated from mosquitoes dissected from an independent cage. Statistical significance calculated by Kruskal-Wallis with Dunn’s multiple comparisons test. Black lines represent mean while the error bars represent SEM. C. Images from time lapse recording showing two ‘waving-flipping’ *trp1Δ14* sporozoites vs normal gliding motility of a wt sporozoite. Red arrows indicate the tip of the sporozoite in each panel. D. Motility patterns observed in various TRP1 C-terminus domain mutant salivary gland sporozoites compared to wild type. ‘n’ equals the total number of sporozoites quantified from each parasite line. The pie chart in green depicts the proportion of moving and non-moving sporozoites overall in each parasite line, where both productive and unproductive motility accounts for the ‘moving’ sporozoites. Attached and floating sporozoites were combined together into the ‘non-moving’ category (Data collected from two independent cage feed experiments). E. Natural transmission assay comparing parasitemia in mice infected with wt and *trp1Δ14* and *trp1Δctd* infected mosquitoes. Vertical arrows indicate the day when parasites are detected in the blood (prepatency). t/n: number of mice showing blood stage infection / number of challenged mice. Black lines represent mean while the error bars represent SEM.

The cytoplasmic tail of *P. falciparum* TRP1 but not of TRAP can complement the cytoplasmic tail of *P. berghei* TRP1.

In contrast to TRAP, the cytoplasmic tail of TRP1 lacks a penultimate tryptophan and shows highly divergent lengths across different *Plasmodium* species (Figure 4A). The cytoplasmic tail of *P. falciparum* TRP1 is only 19 amino acids long compared to 44 amino acids in *P. berghei* and even 97 amino acids in *P. vivax*. ColabFold predictions of the cytoplasmic tails fused to the TMD of TRP1 reveal a similar α-helical region next to the transmembrane domain followed by an unstructured C-terminal segment in *Pb*TRP1 (Figure 4B) that is all but absent in *Pf*TRP1. In contrast, *Pb*TRAP exhibits a reduced α-helical region and an extended unstructured segment that folds back toward the transmembrane domain (Figure 4B). Considering the striking differences in motility in parasite lines lacking the 14 or 19 amino acids from the *P. berghei* TRP1 C-terminus, we next probed if the shorter *P. falciparum* tail or the cytoplasmic tail of *P. berghei* TRAP could complement the one of TRP1 by generating lines of *P. berghei* expressing TRP1 with the C-terminus of *Pb*TRAP (*Pbtrap-swap*) or of *Pf*TRP1 (*Pftrp1-swap*) (Supplementary Figure 5). As expected, the resulting parasite lines infected mosquitoes at rates comparable to wild type parasites (Supplementary Figure 5). Yet, only few parasites expressing TRP1 with the cytoplasmic tail of TRAP did proceed to the salivary glands, while those expressing the short cytoplasmic tail of *P. falciparum* TRP1 readily did so (Figure 4C).

**Figure 4.**
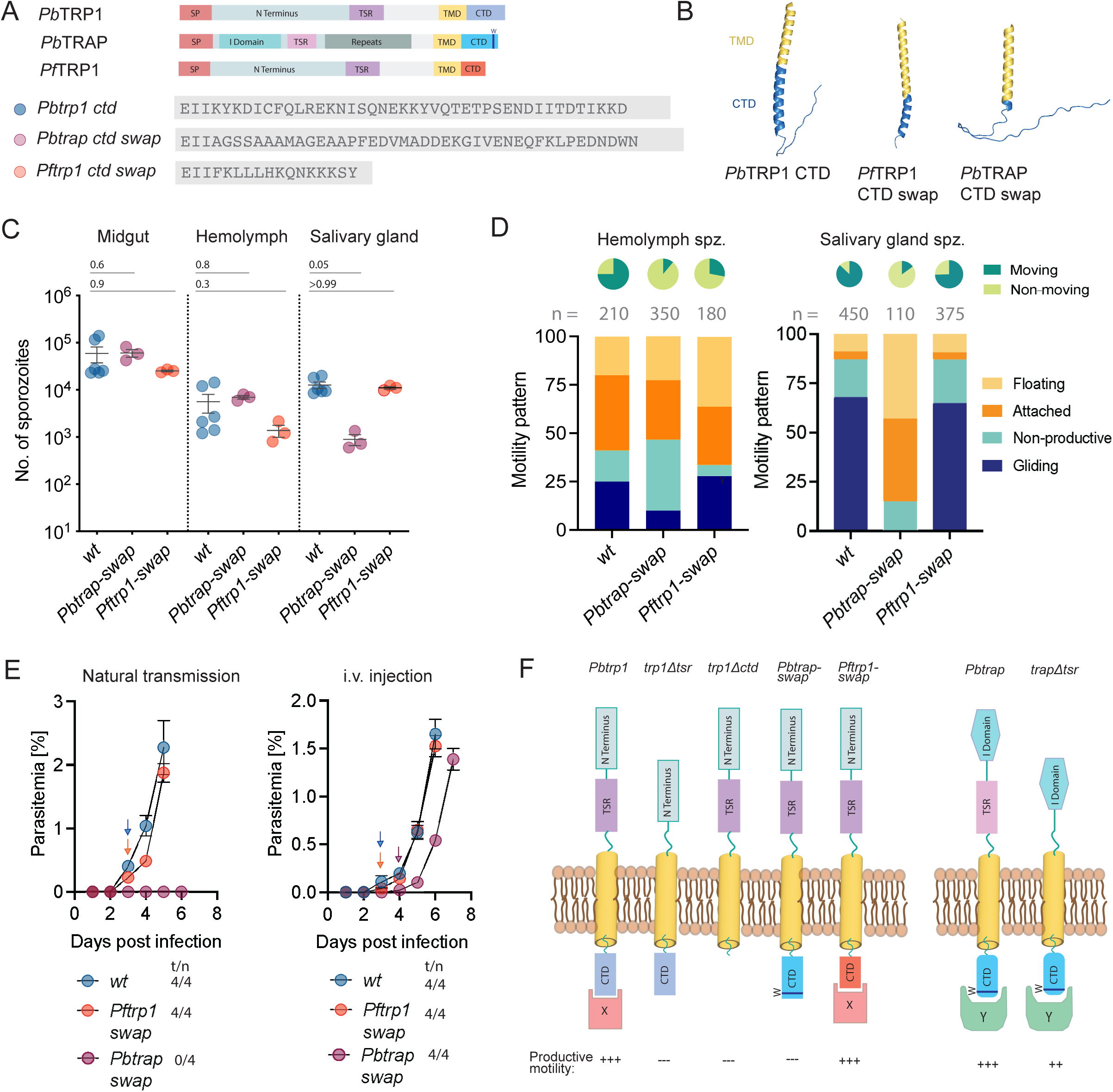
The cytoplasmic tail of *P. falciparum* TRP1 but not of TRAP can complement the cytoplasmic tail of *P. berghei* TRP1. B. Schematic representation of the indicated TRP1 C-terminus domain swap mutants and their amino acid sequences. C. ColabFold prediction of C-terminal domains (blue) of *Pb*TRP1, *Pb*TRAP and *Pf*TRP1. For predictions the C-terminal domains were ‘fused’ to the transmembrane domain (TMD) of *Pb*TRP1 (yellow). D. Total numbers of sporozoites per midgut, hemolymph and salivary gland of various C-terminal deletion mutants; Each dot represents an average number of sporozoites calculated from mosquitoes dissected from an independent cage feed. t/n: number of mice showing blood stage infection / number of challenged mice. Statistical significance calculated by Kruskal-Wallis with Dunn’s multiple comparisons test (Data from three independent cage feeds). Black lines represent mean while the error bars represent SEM. E. Motility patterns of wt and TRP1 C-terminal domain swap mutant hemolymph and salivary gland sporozoites. ‘n’ equals the total number of sporozoites quantified from each parasite line. The pie chart in green depicts the proportion of moving and non-moving sporozoites overall in each parasite line, where both productive and unproductive motility accounts for the ‘moving’ sporozoites. Attached and floating sporozoites were combined together into the ‘non-moving’ category. F. Natural transmission and i.v. injection assays comparing parasitemia in mice infected with *Pbtrap swap*, *Pftrp1 swap* or *wt* infected mosquitoes. t/n: number of mice showing blood stage infection / number of challenged mice. Black lines represent mean while the error bars represent SEM. Black lines represent mean while the error bars represent SEM. G. Summary cartoon suggesting the binding of different cytoplasmic connectors to TRP1 and TRAP and the role of the different protein domains in motility.

Investigating the movement patterns of hemolymph and salivary gland derived sporozoites showed some gliding movement in hemolymph-derived *Pbtrap-swap* parasites (i.e. expressing TRP1 with the cytoplasmic tail of TRAP) but none in salivary gland derived ones (Figure 4D). In contrast, the parasites expressing TRP1 with the cytoplasmic tail of *P. falciparum* TRP1 moved in a similar fashion to wild type (Figure 4D). Transmission experiments further showed that parasites expressing the cytoplasmic tail of TRAP failed to be transmitted to mice, while those expressing the cytoplasmic tail of *P. falciparum* TRP1 infected mice as efficiently as wild type parasites (Figure 4E). Only i.v. injection of 1,000 sporozoites showed that also sporozoites expressing TRP1 with the cytoplasmic tail of TRAP could infect mice, albeit with an average delay of 1.5 days, reflecting a 95% decrease in infectivity (25) (Figure 4E, Table 2). Taken together these data indicate that the diverse cytoplasmic tails from *P. falciparum* and *P. berghei* TRP1 are interchangeable and likely link to the same cytoplasmic protein to enable gliding motility. Interestingly, these data also suggest that this cytoplasmic protein is different from the one interacting with TRAP, thus potentially revealing two pathways of adhesin to actin filament linkage that are equally important for migration.

## Discussion

Gliding motility of *Plasmodium* sporozoites is essential for egress from oocysts, invasion of and dispersion within salivary glands, migration in the skin and entry of the liver. Gliding is powered by an actin-myosin motor (1) and depends on proteins on the sporozoite plasma membrane. Several of these proteins are modulating sporozoite motility at specific steps such as TREP for salivary gland entry (21, 26), TLP for skin migration (27, 28) and CSP for liver entry (29, 30). Others are essential for multiple steps such as TRAP, which is important for all steps post oocyst egress (3) and TRP1, which is important for oocyst egress and potentially for salivary gland invasion (14). Yet, whether or how these proteins interact with each other, how they signal or link to the actin filaments is largely not known (5, 26, 31). Also, the different domains of TRAP have been functionally dissected by mutations in the I-domain, the TSR and the cytoplasmic tail, which revealed a model whereby interactions of ligands with the I-domain leads to an outside-in signal that mediates changes in binding of the cytoplasmic tail to actin filaments (11, 12, 21, 22). Here, we compare the role of the sole TSR domains in TRP1 and TRAP and examine the role of the cytoplasmic tail of TRP1 in gliding motility and life cycle progression.

Deletion of the TSR of TRAP led to only a mild reduction in gliding motility and no impairment of life cycle progression, similar to previous findings using point mutations (21). In contrast, deletion or point mutations in the TSR of TRP1 or tagging of TRP1 at the TSR with GFP led to a sharp drop in the capacity of the mutant parasites to invade salivary glands. Parasites lacking the TRP1-TSR were not able to move, revealing a role of the TSR in gliding motility. How this role is mediated, however, remains obscure. TRP1 is suggested to be cleaved between the TSR and the TMD and hence the N-terminal part of TRP1 including the TSR might not be associated with the rest of the protein on the sporozoite surface (14). Yet, deletion of the TSR has dramatic implications. One explanation could be that deletion or mutation of the TSR affects protein levels due to the lack of domain stabilization through glycosylation as was shown for TRAP (32, 33). Despite several attempts we failed to get a clear GFP fusion protein band of the *trp1-gfp-tsr* parasite with western blotting, suggesting that the protein could be degraded. We also failed so far to raise anti-TRP1 antibodies, which would be valuable tools. Hence future work should address the exact timing of cleavage possibly using smaller tags integrated at different parts of the protein. While a rhomboid protease cleaves TRAP (15, 34) the protease processing TRP1 remains unknown.

Mutations of the cytoplasmic tail of TRP1 led to several interesting observations. Firstly, full deletion of the cytoplasmic tail reveals a deficit in gliding motility after entry into the salivary gland, implying TRP1 functions in sporozoite migration. The newly observed pattern of movement termed wave-flipping suggests that in these mutants the parasites lose their contact to the substrate and hence fail to maintain persistent gliding motility *in vitro*. Whether this translates to a gliding defect in 3D or *in vivo* remains to be demonstrated. Secondly, replacement of the 44 amino acid long tail by the much shorter 19 amino acid long tail of *P. falciparum* TRP1 yielded fully functional sporozoites. Previous work also showed that the C-terminus of *Pb*TRAP could be replaced with the C-terminus of *Pf*TRAP or the C-terminus of *Toxoplasma gondii* MIC2 (23, 35) and even with the C-terminus of TLP (35) without much loss of function. In contrast, the replacement of the TRP1 C-terminus with the cytoplasmic tail of TRAP led to sporozoites that could barely enter salivary glands and the few that managed, failed to undergo gliding motility. This suggests that the two proteins bind to two different partners with their cytosolic domains (Figure 4F). Possibly these different proteins play a role in linking to and/or organizing actin filaments to allow their proper myosin-driven retrograde flow and hence force coupling to the substrate. Clearly, the identification of binding partners of TRAP and TRP1 will be essential to improve our knowledge on gliding motility. Whether ligand binding to the TRP1-TSR and possible outside-in signaling plays a role in gliding remains to be investigated in tight connection to investigations of TRP1 cleavage.

Localization of proteins on the sporozoite surface is notoriously difficult as many of them show strong localization to internal vesicles or the ER (16, 17). Despite several tagged parasite lines expressing TRP1 with GFP, the localization of TRP1 is still enigmatic. Internal tagging of TRP1 with GFP at the cytosolic face of the TMD yielded functional sporozoites with no defects in life cycle progression. This suggests that the fusion protein functions like the endogenous TRP1. It also suggests that the cytoplasmic tail can function even if placed at some distance to the plasma membrane. The curious static localization of TRP1-TMD-GFP in different types of patterns in different sporozoites also raises a number of questions. Is the TRP1 function localized to these hotspots? How does a protein fixed on the surface engage with actin filaments that are transported at great speed towards the end? Could TRP1 function as a modulator of TRAP, e.g. by dissociating TRAP from actin filaments? How is TRP1 trafficked to the surface? While TRAP is accumulated in micronemes and secreted at the front, the localization of TRP1 to micronemes is less evident. Some signals can be observed in a region that is full of micronemes, but it is weaker than the patchy signal near the surface. Clearly more work is needed to understand the intricacies of the different sporozoite surface proteins and how they interact with the substrate on the outside and actin filaments on the insight.

In conclusion, our data show that TRP1 is an essential protein for sporozoite salivary gland invasion, gliding motility and transmission of malaria that acts in a way that is different from TRAP function. A better understanding of the functions of sporozoite surface proteins will not only improve our knowledge of gliding motility and malaria transmission but might help in improving existing vaccines.

## Materials and Methods

### Ethics statement

Animal experiments were performed according to guidelines of the Federation of European Laboratory Animal Science Associations and the Society of Laboratory Animal Science and approved by the responsible German authorities (Regierungspräsidium Karlsruhe). Mice were obtained from Janvier or Charles River Laboratories and kept in the dedicated animal facility of Heidelberg University according to current guidelines (3 mice per cage, ad libitum food and water, environmentally enriched cages).

### Animals and Parasites

Female 4 to 6-week-old Swiss CD1 mice from Janvier were used to generate mutant *P. berghei* parasite lines based on the *P. berghei* strain ANKA and to propagate parasites and to generate gametocytes or to infect mosquitoes for sporozoite production. For transmission experiments, 4 to 6-week-old C57/Bl6 mice from Charles River laboratories were used. *A. stephensi* mosquitoes were reared and maintained in our insectary according to standard procedures.

### Cloning of Plasmid Construct for Parasite Transfection

For the generation of TRP1 (PBANKA_0707900) C-terminal and TSR domain mutants, the Pb238 vector (36) was used as the backbone. This vector contains the human *dhfr* selection cassette under the control of the ef1α promoter and a *P. berghei dhfr* terminator. All constructs were generated by Gibson assembly and transfected into wild-type parasites as SacII/XhoI-digested linear fragments for integration via double homologous recombination. Clonal isogenic lines were obtained by pyrimethamine selection followed by limiting dilution. For internal GFP tagging, a 767 bp eGFP fragment flanked by glycine linkers was inserted using primer pairs P19-P20 or P21-P22 respectively, either upstream of the TSR domain (*trp1-gfp-tsr*) using primer pairs P15-P16 and P17-P18, or before the C-terminus (*trp1-tmd-gfp*) using primer pairs P23-24 and P17-25 via a four-fragment Gibson assembly. The *trp1Δtsr* deletion mutant was generated by assembling fragments spanning the *trp1* ORF upstream and downstream of the TSR domain using primer pairs P23-28, P17-29. Point mutations in conserved tryptophan residues (W591A, W594A) were introduced by assembling overlapping PCR fragments using primer pairs P23-26, P17-27 carrying the desired substitutions (*trp1-tsr-AA*). The *trp1-tsr^TRAP^* line was generated similarly, using primer pairs P23-30, P17-31 replacing the TSR domain with that of TRAP amplified from wild-type genomic DNA. All PCR fragments were assembled using Gibson assembly.

C-terminal deletion mutants were generated by cloning truncated *trp1* ORFs into the TRP1 C-terminal GFP vector (14) using SacII and BamHI, replacing the *gfp* cassette with an *hdhfr-yfcu* selection marker using primer pairs P32-33 (*trp1Δ3*), P32-34 (*trp1Δ14*), P32-35 (*trp1Δ19*). Homologous regions comprised ∼1 kb upstream of the deletion site and the terminal 609 bp of the C-terminus plus ∼500 bp downstream sequence, preserving the adjacent gene PbANKA_070800 due to the short intergenic distance (291 bp).

For domain swap mutants, the TRP1 C-terminal GFP vector backbone was used to replace the Pbtrp1 C-terminus with either the 98 bp *Pftrp1* C-terminal region using primer pairs P36-41, P42-43, P17-44 or the 178 bp *Pbtrap* C-terminal region using primer pairs P36-37, P38-39, P17-40, together with the *Pbdhfr* 3′ UTR. These constructs excluded the *gfp* coding sequence and were assembled using Gibson assembly.

For generating *trap* tsr domain mutant PlasmoGem vector (*Pb*GEM-107890) (36) was used as a backbone. The DNA sequence encoding the TSR region (C238–P281; 44 amino acids) was deleted by site-directed mutagenesis using primers P45 and P46. The mutated *trap* coding sequence, including its 5′ and 3′ UTRs, was subsequently excised (SacII and EcoRV) from the pGEM-TRAPΔTSR construct and cloned into the *Pb*238 intermediate vector, which already contained the trap 3′ UTR downstream of the selection cassette. The final construct was digested with SacII and KpnI, purified, and transfected into wild-type parasites for genomic integration. Parasites were selected using pyrimethamine post transfection. Isogenic parasite lines were generated using limiting dilution.

### Generation of *P. berghei* Parasite Lines

TRP1 and TRAP C-terminus domain and TSR domain mutants of *P. berghei* were generated through double homologous recombination (Supplementary Figure 1-5). To achieve this, linearized plasmid constructs were transfected into the parental *P. berghei* strains—PbANKA wild type for TRP1 and TRAP using standard protocols (37). Transgenic parasites were selected via pyrimethamine (0.07 mg/mL) administered in the drinking water of infected mice. Clonal parasite lines were established from mixed populations by limiting dilution, where a single blood-stage parasite was intravenously injected into 8-9 Swiss mice.

Once parasitemia reached 1–3%, blood was collected by cardiac puncture under anesthesia (120 mg/kg ketamine and 16 mg/kg xylazine, intraperitoneally). Cryopreserved stabilates were prepared and stored in liquid nitrogen. For genotyping, infected blood was lysed using 0.093% saponin, and parasites were resuspended in 200 µL PBS. Genomic DNA was extracted with the Qiagen Blood and Tissue Kit according to the manufacturer’s instructions. All resulting isogenic lines were validated by genotyping PCR, using primers listed in Supplementary Table 1. Lines with the correct genotype were considered isogenic.

### Mosquito Infection

Donor mice were intraperitoneally infected with the appropriate *Plasmodium* stabilates to generate infections for mosquito feeding. Parasites were expanded for 4–5 days until parasitemia reached ∼2%, after which donor mice were bled by cardiac puncture. Fresh blood was used to transfer 2 × 10^7^ parasites into 2 naïve recipient mice. These mice were maintained for an additional 3–4 days, depending on gametocyte development, which was assessed by exflagellation assays. Mice exhibiting sufficient gametocytemia (≥3–5 exflagellation events per microscopic field) were used for mosquito feeding (Around 300 female *Anopheles stephensi* mosquitoes). Prior to feeding, mice were anesthetized with ketamine (100 mg/kg) and xylazine (3 mg/mL) and placed on top of mosquito cages under dimmed light conditions. Mosquitoes were allowed to feed for 20–30 min, with mice repositioned every 10 min to ensure uniform access. Following feeding, mosquitoes were maintained at 21°C and 80% relative humidity and provided with both humid sugar and salt pads for optimal survival.

### Analysis of Oocyst Development

To assess mosquito midgut infection rates, midguts were dissected 10–14 days post-infection and stained with mercurochrome to visualize *Plasmodium* oocysts. Following dissection in 100 µL PBS, midguts were permeabilized in 1% NP-40 (in PBS) for 30 min. Samples were then incubated in 0.1% mercurochrome (in PBS) for at least 1 h to enhance contrast between the oocyst wall and the surrounding midgut musculature. After staining, midguts were washed 2–3 times with PBS until the wash solution was clear. Stained midguts were transferred onto glass slides using a Pasteur pipette, covered with a coverslip, and sealed with wax. Imaging was performed using an Axiovert 200M fluorescence microscope (Carl Zeiss) with a 10X objective.

### Sporozoite Isolation and Counting

Sporozoites were isolated from mosquito midguts, hemolymph, and salivary glands between days 11 and 24 post-infection, depending on the experimental objective. Midgut sporozoites were harvested between days 10 and 14, hemolymph sporozoites between days 14 and 16, and salivary gland sporozoites between days 17 and 24 post-infection. For parasite lines exhibiting defects in oocyst egress or salivary gland invasion, sporozoite numbers were quantified at multiple time points (days 14, 17/18, 20, and 22 post-infection) to validate the phenotype. For sporozoite quantification, midguts or salivary glands from at least 10 mosquitoes were dissected in PBS or RPMI medium, homogenized using a pestle, and free sporozoites were counted using a Neubauer hemocytometer. Midgut-derived sporozoites were diluted 1:10 prior to counting to facilitate accurate counting. Samples were allowed to settle for 5 min before counting using a phase-contrast light microscope (Carl Zeiss) with a 40X objective. Hemolymph sporozoites were isolated by anesthetizing mosquitoes on ice for at least 30 min, followed by removal of the terminal abdominal segment using a fine needle. Mosquitoes were flushed by injecting chilled PBS into the thorax with a Pasteur pipette, allowing hemolymph to drain from the abdomen. Collected hemolymph was transferred into 1.5-mL microcentrifuge tubes (Eppendorf) and centrifuged at 11,000 rpm for 3 min. The supernatant was discarded, and the pellet was resuspended in 100 µL fresh PBS. Sporozoite counts were performed as described for midgut and salivary gland samples. For each replicate, a minimum of 20 female mosquitoes were processed.

### Sporozoite Motility Assay

Sporozoite gliding motility assays were performed using salivary gland– or hemolymph-derived sporozoites. Salivary glands from 20–30 infected mosquitoes were dissected in 50 µL ice-cold RPMI and homogenized with a pestle to release sporozoites from salivary glands. Sporozoites were purified by density gradient centrifugation using 17% Accudenz without brakes. Purified pellets were resuspended in 200 µL RPMI supplemented with 3% bovine serum albumin (BSA) and transferred to uncoated, optical-bottom 96-well plates. Hemolymph sporozoites were isolated from approximately 20 infected mosquitoes as described above, centrifuged for 3 min at 13.000 rpm at room temperature, and resuspended in 200 µL RPMI or PBS supplemented with 3% BSA before transfer to 96-well plates. Independent of sporozoite origin, plates were centrifuged for 3 min at 1.500 rpm and immediately imaged. Time-lapse imaging was performed using an Axiovert 200M fluorescence microscope (Carl Zeiss). Movies were acquired in differential interference contrast (DIC) mode with a 25X objective at one frame every 3 seconds and analyzed using FIJI and categorized into five patterns as previously described (18): Gliding, Floating, Attached, Non-productive and a newly observed motility pattern that was described as Wave-Flipping.

### Imaging of parasite lines

Oocysts and salivary gland sporozoites were imaged at 11–14 and 17–24 days post-infection, respectively. Midguts or salivary glands were dissected, mounted in RPMI or PBS containing Hoechst 33342 (1:1000 dilution from a 10 mg/mL DMSO stock), sealed, and imaged immediately using a spinning disc confocal microscope (PerkinElmer) at 100× magnification (Frénal et al., 2017a). For isolated sporozoites, parasites were collected at 17–21 days post-infection, transferred to 8-well optical-bottom plates (Ibidi), mixed 1:1 with RPMI containing 6% BSA and Hoechst 33342 (1:1000), centrifuged (3 min, 1500 rpm; Heraeus Multifuge S1), and imaged immediately at 100× magnification. For immunofluorescence analysis of sporozoites, infected midguts or salivary glands were dissected in ice-cold PBS or RPMI (Eppendorf tubes). Sporozoites were mechanically released, purified by density gradient centrifugation, resuspended in PBS, and seeded onto 8-well optical-bottom plates (Ibidi). Parasites were activated with an equal volume of PBS containing 6% BSA, centrifuged (3 min, 13,000 rpm, RT), and allowed to glide for 20–60 min at room temperature. Samples were fixed with 4% PFA in PBS (1 h, RT), washed three times with PBS, and blocked (PBS + 2% BSA) or blocked/permeabilized (PBS + 2% BSA + 0.5% Triton X-100) for 1 h at RT or overnight at 4°C. Primary antibodies were incubated for 1 h at RT or overnight at 4°C, followed by three PBS washes and 1 h incubation with secondary antibodies (RT, dark). After final washes, samples were maintained in PBS and imaged immediately or stored at 4°C until use.

### Transmission to Mice

To assess the transmission potential of the generated parasite lines, mice were infected either by natural transmission via mosquito bite or by intravenous injection of salivary gland–derived sporozoites. For natural transmission, mosquitoes infected 17–24 days earlier were distributed into cups containing 10 mosquitoes each and starved for 6–8 hours prior to feeding. Naïve C57BL/6 mice were anesthetized by intraperitoneal injection of ketamine (87.5 mg/kg) and xylazine (3 mg/mL) and placed ventral side down on the mosquito cups for approximately 20-30 min. Mosquitoes that successfully blood-fed were dissected immediately or the following day to quantify salivary gland sporozoite loads. For intravenous sporozoite challenge, salivary glands were dissected from mosquitoes infected 17–24 days earlier in PBS and sporozoites were released and diluted to a final concentration of 1000 sporozoites per 100 µL. Freshly prepared sporozoite suspensions were injected into the tail vein of naïve C57BL/6 mice. Parasitemia was monitored daily from days 3 to 20 post-infection by examination of Giemsa-stained blood smears using a light microscope (Carl Zeiss) equipped with a counting grid, and survival was monitored for up to 30 days. The time from infection to the first detectable blood-stage parasitemia was defined as the prepatent period.

## Acknowledgements

We thank Miriam Reinig and many students for help in mosquito rearing and Justin Boddey, Julia Sattler, Mirko Singer and Kevin Walz for comments on the manuscript and many helpful discussions. MRC and DK were members of the Heidelberg Biosciences International Graduate School (HBIGS). MM was a member of the Molecular Biotechnology Master Program and BL was a member of the Medical Doctoral Program at Heidelberg University. We acknowledge the microscopy support from the Infectious Diseases Imaging Platform (IDIP) at the Center for Integrative Infectious Disease Research. The *Plasmodium* database PlasmoDB facilitated this work.

## Author contribution

MRC and FF designed research; MRC, MM, BL, DK and BJ performed research; MRC, MM, DK, JK analysed data, MRC, DK, JK, JMS, MS, FH and FF supervised parts of the study, FF acquired funding and MRC, FH and FF wrote the paper.

## References

1. K. Frénal, J.-F. Dubremetz, M. Lebrun, D. Soldati-Favre, Gliding motility powers invasion and egress in Apicomplexa. Nat. Rev. Microbiol. 15, 645–660 (2017).

2. F. Frischknecht, K. Matuschewski, Plasmodium Sporozoite Biology. Cold Spring Harb. Perspect. Med. 7, a025478 (2017).

3. A. A. Sultan, et al., TRAP Is Necessary for Gliding Motility and Infectivity of Plasmodium Sporozoites. Cell 90, 511–522 (1997).

4. B. J. Morahan, L. Wang, R. L. Coppel, No TRAP, no invasion. Trends Parasitol. 25, 77–84 (2009).

5. K. Beyer, et al., Limited Plasmodium sporozoite gliding motility in the absence of TRAP family adhesins. Malar. J. 20, 430 (2021).

6. R. G. Douglas, M. Reinig, M. Neale, F. Frischknecht, Screening for potential prophylactics targeting sporozoite motility through the skin. Malar. J. 17, 319 (2018).

7. D. Jacot, et al., An Apicomplexan Actin-Binding Protein Serves as a Connector and Lipid Sensor to Coordinate Motility and Invasion. Cell Host Microbe 20, 731–743 (2016).

8. J. F. Stortz, et al., Formin-2 drives polymerisation of actin filaments enabling segregation of apicoplasts and cytokinesis in Plasmodium falciparum. eLife 8, e49030 (2019).

9. K. A. Quadt, M. Streichfuss, C. A. Moreau, J. P. Spatz, F. Frischknecht, Coupling of Retrograde Flow to Force Production During Malaria Parasite Migration. ACS Nano 10, 2091–2102 (2016).

10. D. Klug, et al., Evolutionarily distant I domains can functionally replace the essential ligand-binding domain of Plasmodium TRAP. eLife 9, e57572 (2020).

11. T. A. Springer, M. L. Dustin, Integrin inside-out signaling and the immunological synapse. Curr. Opin. Cell Biol. 24, 107–115 (2012).

12. F. Braumann, et al., Conformational change of *Plasmodium* TRAP is essential for sporozoite migration and transmission. EMBO Rep. 24, e57064 (2023).

13. K. E. Swearingen, et al., Interrogating the Plasmodium Sporozoite Surface: Identification of Surface-Exposed Proteins and Demonstration of Glycosylation on CSP and TRAP by Mass Spectrometry-Based Proteomics. PLOS Pathog. 12, e1005606 (2016).

14. D. Klug, F. Frischknecht, Motility precedes egress of malaria parasites from oocysts. eLife 6, e19157 (2017).

15. I. Ejigiri, et al., Shedding of TRAP by a Rhomboid Protease from the Malaria Sporozoite Surface Is Essential for Gliding Motility and Sporozoite Infectivity. PLoS Pathog. 8, e1002725 (2012).

16. A. F. Carey, et al., Calcium dynamics of Plasmodium berghei sporozoite motility. Cell. Microbiol. 16, 768–783 (2014).

17. J. Kehrer, F. Frischknecht, G. R. Mair, Proteomic Analysis of the Plasmodium berghei Gametocyte Egressome and Vesicular bioID of Osmiophilic Body Proteins Identifies Merozoite TRAP-like Protein (MTRAP) as an Essential Factor for Parasite Transmission. Mol. Cell. Proteomics 15, 2852–2862 (2016).

18. C. Thieleke-Matos, K. Walz, F. Frischknecht, M. Singer, Overcoming the egress block of Plasmodium sporozoites expressing fluorescently tagged circumsporozoite protein. Mol. Microbiol. 121, 565–577 (2024).

19. M. Singer, F. Frischknecht, Fluorescent tagging of *Plasmodium* circumsporozoite protein allows imaging of sporozoite formation but blocks egress from oocysts. Cell. Microbiol. 23 (2021).

20. J. Kehrer, et al., A Putative Small Solute Transporter Is Responsible for the Secretion of G377 and TRAP-Containing Secretory Vesicles during Plasmodium Gamete Egress and Sporozoite Motility. PLOS Pathog. 12, e1005734 (2016).

21. K. Matuschewski, Plasmodium sporozoite invasion into insect and mammalian cells is directed by the same dual binding system. EMBO J. 21, 1597–1606 (2002).

22. K. Wengelnik, et al., The A-domain and the thrombospondin-related motif of Plasmodium falciparum TRAP are implicated in the invasion process of mosquito salivary glands. EMBO J. 18, 5195–5204 (1999).

23. S. Kappe, et al., Conservation of a gliding motility and cell invasion machinery in Apicomplexan parasites. J. Cell Biol. 147, 937–944 (1999).

24. S. Hegge, M. Kudryashev, A. Smith, F. Frischknecht, Automated classification of *Plasmodium* sporozoite movement patterns reveals a shift towards productive motility during salivary gland infection. Biotechnol. J. 4, 903–913 (2009).

25. J. P. Vanderberg, Development of infectivity by the Plasmodium berghei sporozoite. J. Parasitol. 61, 43–50 (1975).

26. S. Hegge, et al., Direct manipulation of malaria parasites with optical tweezers reveals distinct functions of Plasmodium surface proteins. ACS Nano 6, 4648–4662 (2012).

27. J. K. Hellmann, et al., Environmental Constraints Guide Migration of Malaria Parasites during Transmission. PLoS Pathog. 7, e1002080 (2011).

28. I. Ejigiri, P. Sinnis, Plasmodium sporozoite-host interactions from the dermis to the hepatocyte. Curr. Opin. Microbiol. 12, 401–407 (2009).

29. A. Coppi, et al., Heparan Sulfate Proteoglycans Provide a Signal to Plasmodium Sporozoites to Stop Migrating and Productively Invade Host Cells. Cell Host Microbe 2, 316–327 (2007).

30. A. E. Balaban, et al., The Plasmodium CSP repeats have elastic properties with a critical role in sporozoite motility. EMBO J. 44, 6253–6272 (2025).

31. M. S. Paoletta, S. E. Wilkowsky, Thrombospondin Related Anonymous Protein Superfamily in Vector-Borne Apicomplexans: The Parasite’s Toolkit for Cell Invasion. Front. Cell. Infect. Microbiol. 12, 831592 (2022).

32. S. Lopaticki, et al., Protein O-fucosylation in Plasmodium falciparum ensures efficient infection of mosquito and vertebrate hosts. Nat. Commun. 8, 561 (2017).

33. S. Lopaticki, et al., Tryptophan C-mannosylation is critical for Plasmodium falciparum transmission. Nat. Commun. 13, 4400 (2022).

34. M. Freeman, Rhomboid Proteases and their Biological Functions. Annu. Rev. Genet. 42, 191–210 (2008).

35. K. Heiss, et al., Functional characterization of a redundant Plasmodium TRAP family invasin, TRAP-like protein, by aldolase binding and a genetic complementation test. Eukaryot. Cell 7, 1062–1070 (2008).

36. C. J. Janse, J. Ramesar, A. P. Waters, High-efficiency transfection and drug selection of genetically transformed blood stages of the rodent malaria parasite Plasmodium berghei. Nat. Protoc. 1, 346–356 (2006).

37. C. J. Janse, et al., Plasmodium berghei: Gametocyte production, DNA content, and chromosome-size polymorphisms during asexual multiplication in vivo. Exp. Parasitol. 68, 274–282 (1989).

