## Supplemental Figures 1-5 for "Differential signaling by two *Plasmodium* sporozoite adhesins mediates transmission of malaria parasites"

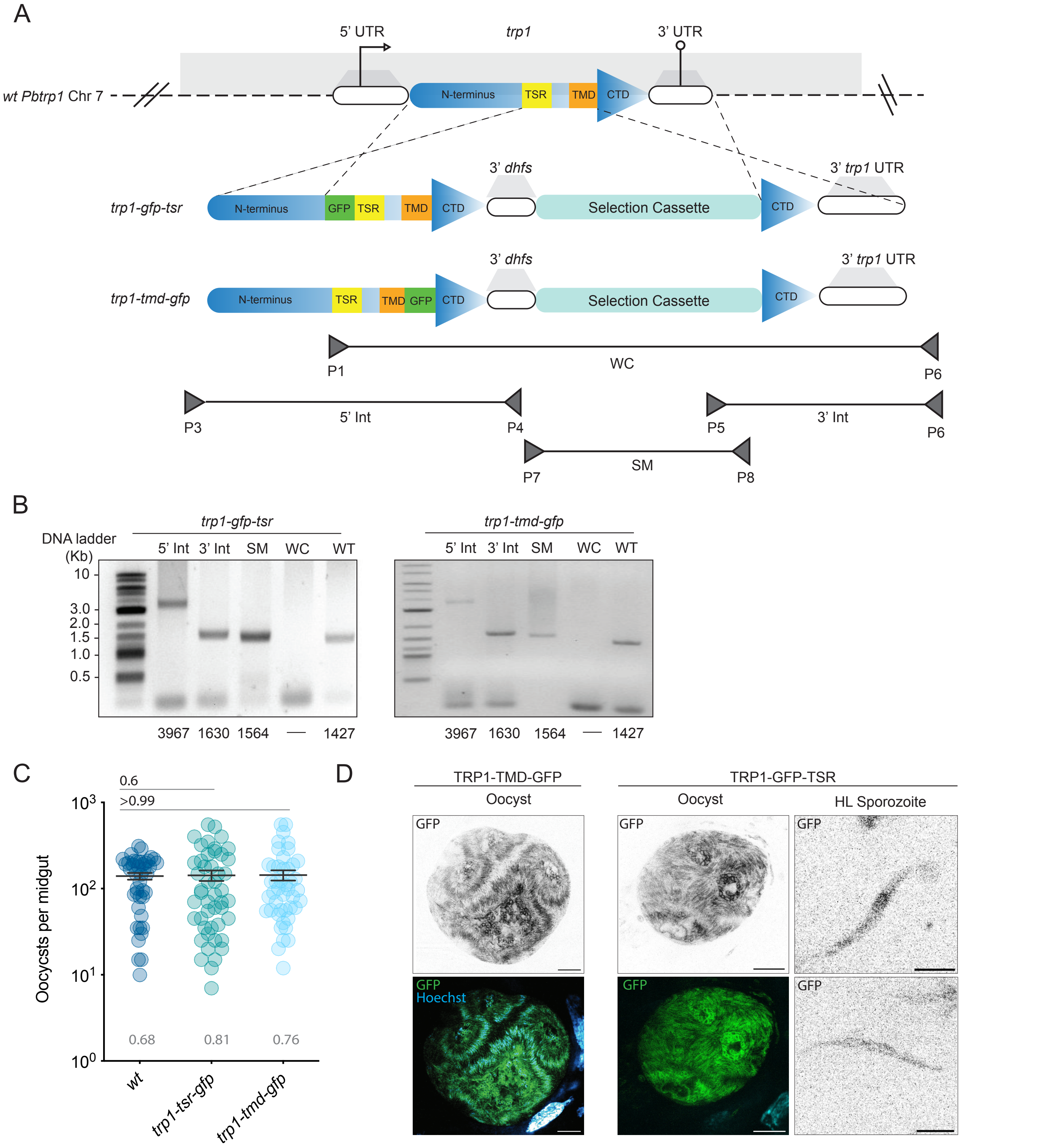


**S1. A.** Generation of TRP1 GFP-tagged mutants. The construct on top indicates the genomic locus and the dotted lines represent double homologous recombination event leading to the generation of GFP tagged mutants. Figures are not drawn to scale; primer placement is approximate. **B.** Genotyping PCRs of *trp1-tmd-gfp* and *trp1-gfp-tsr* parasite lines. Primers used are indicated in scheme S1.A and listed in Table S1. Legends used: Kb: Kilobases, 5'Int: 5' Integration fragment, 3'Int: 3' Integration fragment. SM: Selection marker, WC: Whole construct (Mutant), WT: Wild type control for whole construct.The numbers below indicate the expected band sizes of the PCR products (in bp). **C.** Oocyst numbers per midgut where each dot represents one midgut. Black line indicates the median. Percentage of infection rates are indicated in grey above the X axis. **D.** Localization of GFP in *trp1-tmd-gfp* oocyst (scale bar: 10 µm) compared with Localization of GFP in *trp1-gfp-tsr* oocyst (scale bar: 10 µm) and hemolymph sporozoites (scale bar: 3 µm).

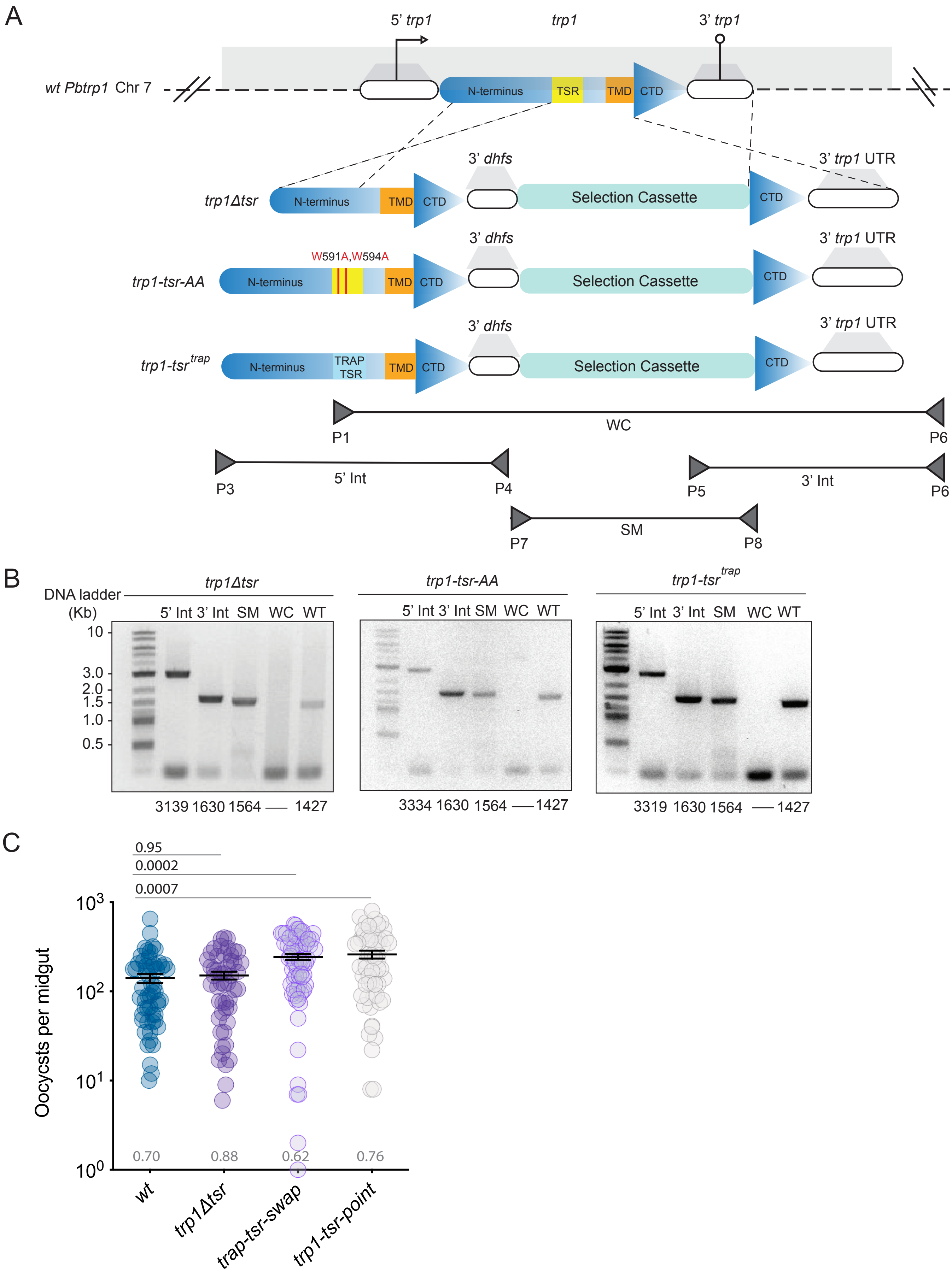


**S2. A.** Generation of TRP1 TSR-mutants. The construct on top indicates the genomic locus and the dotted lines represent double homologous recombination event leading to the generation of TRP1 TSR- mutants. Figures are not drawn to scale; primer placement is approximate. **B.** Genotyping PCRs of TRP1 TSR-mutant parasite lines. Primers used are indicated in scheme S2.A and listed in Table S1. Legends used: Kb: Kilobases, 5'Int: 5' Integration fragment, 3'Int: 3' Integration fragment. SM: Selection marker, WC: Whole construct (Mutant), WT: Wild type control for whole construct.The numbers below indicate the expected band sizes of the PCR products (in bp). **C.** Oocyst numbers per midgut where each dot represents one midgut. Black line indicates the median. Percentage of infection rates are indicated in grey above the X axis.

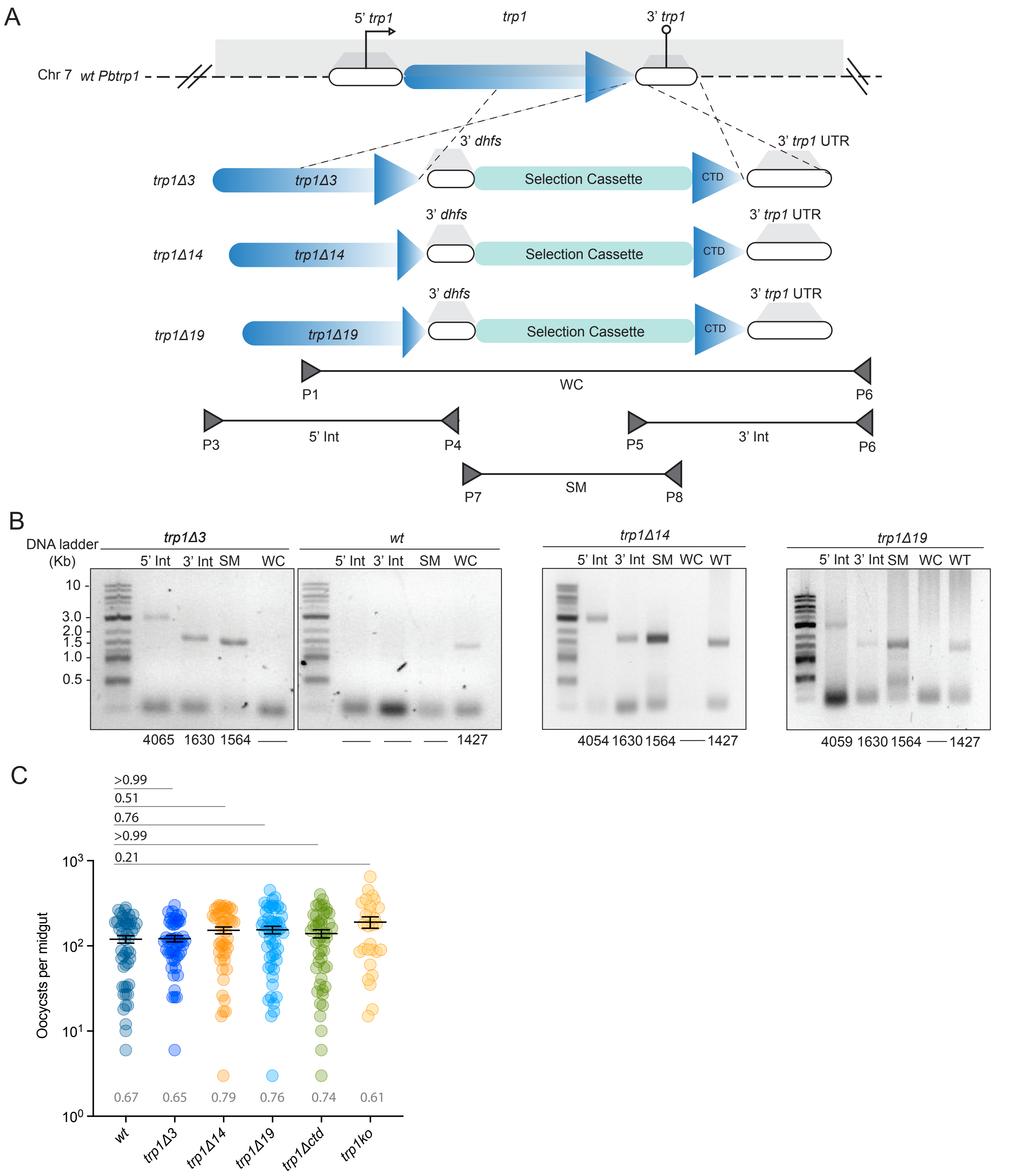


**S3. A.** Generation of TRP1 C-terminus deletion mutants. The construct on top indicates the genomic locus and the dotted lines represent double homologous recombination event leading to the generation of C-terminus deletion mutants. Figures are not drawn to scale; primer placement is approximate. **B.** Genotyping PCRs of C-terminus deletion mutant parasite lines. Primers used are indicated in scheme S3.A and listed in Table S1. Legends used: Kb: Kilobases, 5'Int: 5' Integration fragment, 3'Int: 3' Integration fragment. SM: Selection marker, WC: Whole construct (Mutant), WT: Wild type control for whole construct. The numbers below indicate the expected band sizes of the PCR products (in bp).
**C.** Oocyst numbers per midgut where each dot represents one midgut. Black line indicates the median. Percentage of infection rates are indicated in grey above the X axis.

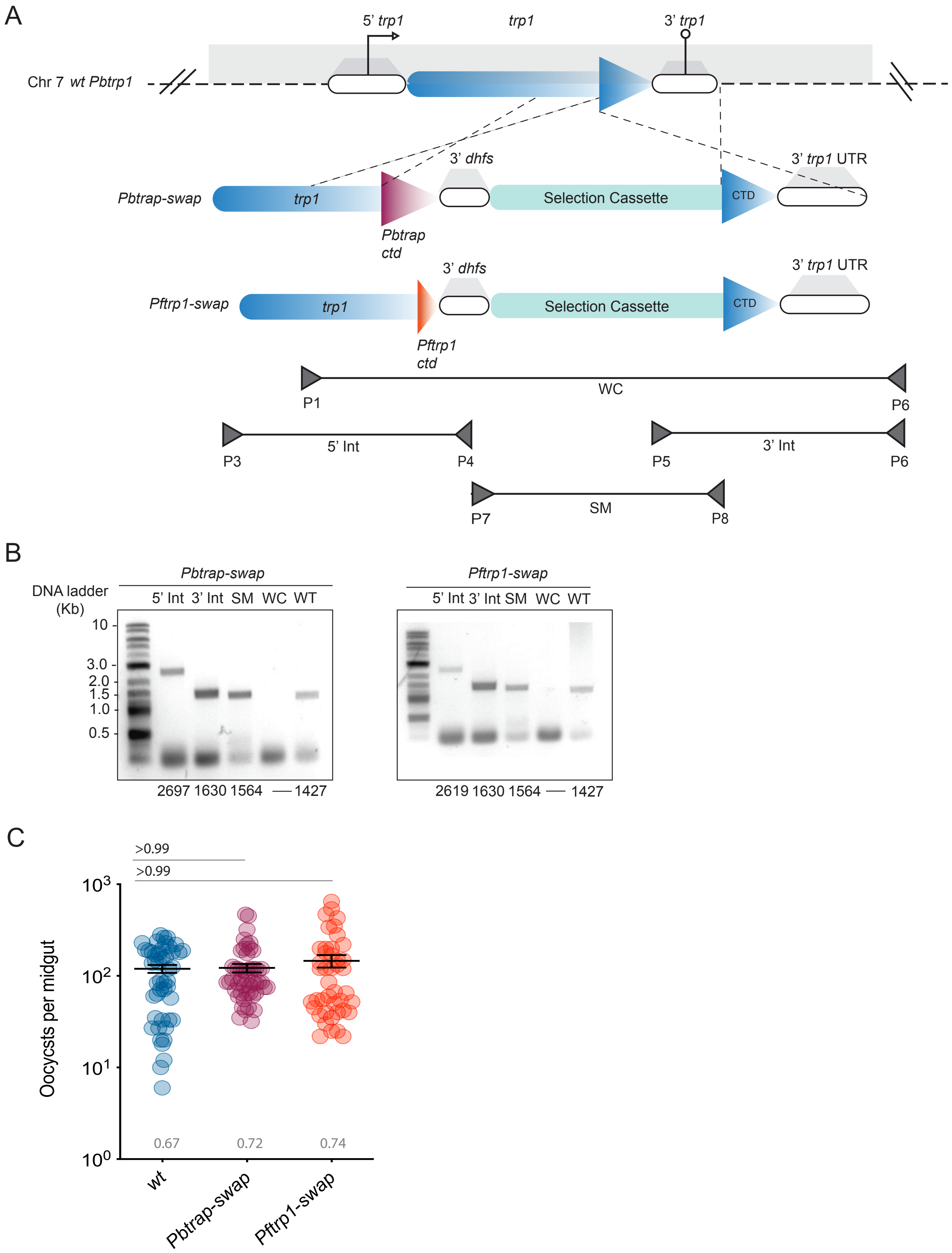


**S4. A.** Generation of TRP1 C-terminus swap mutants. The construct on top indicates the genomic locus and the dotted lines represent double homologous recombination event leading to the generation of C-terminus swap mutants. Figures are not drawn to scale; primer placement is approximate. **B.** Genotyping PCRs of C-terminus swap mutant parasite lines. Primers used are indicated in scheme S4.A and listed in Table S1. Legends used: Kb: Kilobases, 5'Int: 5' Integration fragment, 3'Int: 3' Integration fragment. SM: Selection marker, WC: Whole construct (Mutant), WT: Wild type control for whole construct. The numbers below indicate the expected band sizes of the PCR products (in bp). **C.** Oocyst numbers per midgut where each dot represents one midgut. Black line indicates the median. Percentage of infection rates are indicated in grey above the X axis.

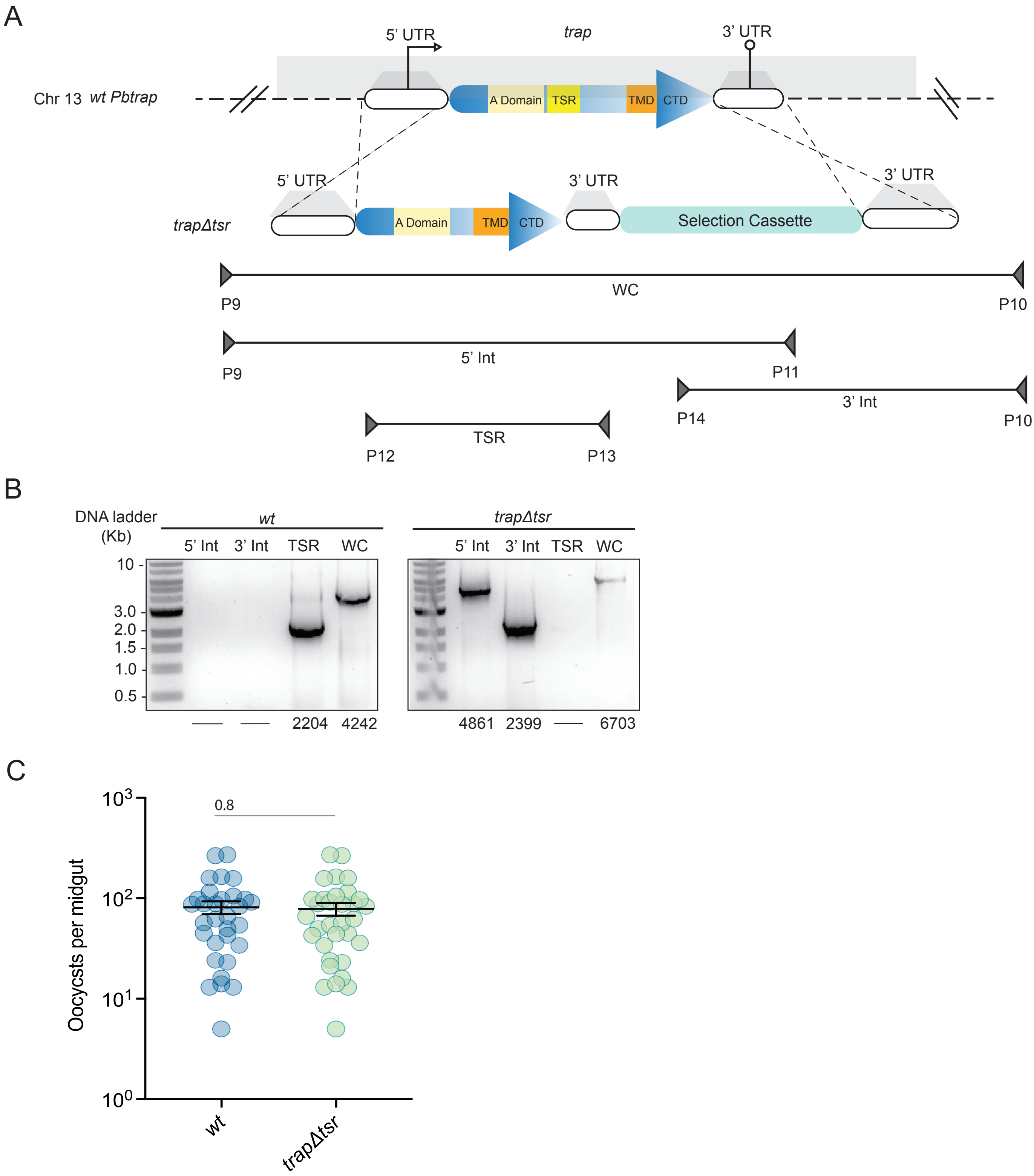


**S5. A.** Generation of *trapΔtsr* parasite line. The construct on top indicates the genomic locus and the dotted lines represent double homologous recombination event leading to the generation of C-terminus swap mutants. Figures are not drawn to scale; primer placement is approximate. **B.** Genotyping PCR of *trapΔtsr* parasite line and corresponding wild type control. Primers used are indicated in scheme S5.A and listed in Table S1. Legends used: Kb: Kilobases, 5'Int: 5' Integration fragment, 3'Int: 3' Integration fragment,WC: Whole construct (Mutant), TSR: TSR domain. The numbers below indicate the expected band sizes of the PCR products (in bp). **C.** Oocyst numbers per midgut where each dot represents one midgut. Black line indicates the median.
