## Supplemental Table 1 for "Differential signaling by two *Plasmodium* sporozoite adhesins mediates transmission of malaria parasites"

**Table S1: Primer list**

| No. | Primer sequence |
|---|---|
| P1 | AATTCCGCGGGATGATAATTGCACTTATTTTGAT |
| P2 | ATTTGGATCCTTATGTTTCAGTTTGTACATATTTTTTTC |
| P3 | ATGTATCGAATTATATTCTTCTTTATTTCATTG |
| P4 | CCCACCGGTGCTTTTTCACGTATATTTTTTTGTTAC |
| P5 | CTTGCACCGGTTTTTATAAAATTTTTATTTATTTATAAGC |
| P6 | GTAGCTCGAGCATCTACTACTCATAATACACTTAGTGGAAGTACG |
| P7 | CCCGCTAGCCCTAGCTAAAAGGTGTGCAAG |
| P8 | CGGGAATTCAAACCATGGTTGGTTCGCT |
| P9 | GAATACATGTAAAAAAGAGAAATTCCTTCG |
| P10 | GTAAAATAAGCGATATAGAAGGGAGC |
| P11 | CGACCGGTAAACTGCATCGTCGCTG |
| P12 | CTGGAAAAAGTTGCTCTTTGTGGAAAATGGGAAG |
| P13 | TAGGATATCCTCCAAACAAAAAATGGACACGTGCAACTA |
| P14 | CTAGCTAGCTTAATCATTCTTCTCATATACTTC |
| P15 | AATCCGCGGATGTATCGAATTATATTCTTCTTTATTTCATTG |
| P16 | TTCTCCTTTACTCATTCCTCCATTACATTTTATTTTTGTATG |
| P17 | TGTATGAAATTACTTTTAAACG |
| P18 | GGAGGATTTAGTTCTTGGTCAGATTG |
| P19 | CATACAAAAATAAAATGTAATGGAGGAATGAGTAAAGGAGAAGAACTTTTC |
| P20 | CAATCTGACCAAGAACTAAATCCTCCTTTGTATAGTTCATCCATGCC |
| P21 | TATCTTGTTTATTTAATTGTCGGAGGATGAGTAAAGGAGAAGAACTTTTC |
| P22 | CTTTATATTTTATAATTTCTTTTCCTtCCTTTGTATAGTTCATCCATGCC |
| P23 | AATGATATACAACTATAGCATGG |
| P24 | TTCTCCTTTACTCATTCCTCCGACAATTAAATAAACAAGAT |

| | |
|---|---|
| P25 | CATGGATGAACTATACAAAGGAGGAAAAGAAATTATAAAATATAAAG |
| P26 | GATTTAGTACATTCTGAGGCATCTGATGCAGAACTAAAATTAC |

| | |
|---|---|
| P27 | CAAAAATAAAATGTAATTTTAGTTCTGCATCAGATGCCTCAGAATGTACTAAAT CATG |
| P28 | GTAACTTCTTTATTTGAATCATTTATATCATTACATTTTATTTTTGTATGTTTACA C |
| P29 | GTGTAAACATACAAAAATAAAATGTAATGATATAAATGATTCAAATAAAGAAGTT AC |
| P30 | CCATTCTTCCCATTTTCCACAATTACATTTTATTTTTGTATGTTTACAC |
| P31 | GTAAGGTTCGTGATTGCCCAGATATAAATGATTCAAATAAAGAAGTTAC |
| P32 | AATTCCGCGGGATGATAATTGCACTTATTTTGAT |
| P33 | ATTTGGATCCTTATATGGTATCTGTAATTATATCATTTTCAG |
| P34 | ATTTGGATCCTTATGTTTCAGTTTGTACATATTTTTTTTC |
| P35 | ATTTGGATCCTTAATATTTTTTTTCATTTTGAG |
| P36 | TAAAATGATATACAACTATAGC |
| P37 | CTACTTCCTGCTATAAAATTTTCTTTGACAATTAAATAAAC |
| P38 | GTTTATTTAATTGTCAAAGAAAATTTTATAGCAGGAAGTAGC |
| P39 | GGGCTTGCACACCTTTTAGCTATTAGTTCCAGTCATTATCTTC |
| P40 | GAAGATAATGACTGGAACTAATAGCTAAAAGGTGTGCAAG |
| P41 | GTAATTTAAAAATATGCCACTTTTCTTTGACAATTAAATAAAC |
| P42 | GTTTATTTAATTGTCAAAGAAAAGTGGCATATTTTTAAATTACTC |
| P43 | GCTTGCACACCTTTTAGCTATTAGTATGATTTTTTTTTGTTTTG |
| P44 | CAAAAAAAAATCATACTAATAGCTAAAAGGTGTGCAAGC |
| P45 | AAGAGCAACTTTTTCTACTTCCTGACAAACTTTAG |
| P46 | CCAAAACCGGTAGCTCCTCCTGTC |
